# The effect of lactational low protein diet on skeletal muscle during adulthood and ageing in male and female mouse offspring

**DOI:** 10.1101/2023.09.07.556663

**Authors:** Moussira Alameddine, Atilla Emre Altinpinar, Ufuk Ersoy, Ioannis Kanakis, Ioanna Myrtziou-Kanaki, Susan E Ozanne, Katarzyna Goljanek-Whysall, Aphrodite Vasilaki

## Abstract

Sarcopenia is characterised by loss of skeletal muscle mass and function associated with a reduction in muscle fibres. External factors, like exercise and diet, can also influence skeletal muscle mass and contribute to muscle fibre loss. Maternal programming refers to the effect of maternal environmental factors such as nutrition that lead to phenotypic changes in the offspring. Maternal malnutrition has been linked to a reduction in body weight and impaired development of skeletal muscle of the offspring; however, there are no studies that reported the long-term effect of maternal low protein diet on the ageing of skeletal muscles. This study aimed to examine how maternal protein deficiency during lactation affects skeletal muscle development and ageing in the offspring. Pups born from normally fed mothers were lactated by low protein fed mothers. Post-weaning, mice were either maintained on a low protein diet (LPD) or switched to normal protein diet (NPD). Pups born from normally fed mothers and maintained on NPD during lactation and afterwards were used as control. In males, the diet mainly affected the size of the myofibres without major effect on fibre number and led to a reduced grip strength of ageing mice (24 months). Female mice had a lower body and muscle weight at weaning but caught up with control mice at 3 months. During ageing, muscle weight, myofibre number and survival rate of female pups were significantly affected. These findings highlight longitudinal animal research for nutritional programming and the importance of sexual dimorphism in response to challenges.

**Highlights:**

- Postnatal low protein diet significantly decreases the survival rate of female but not male mice.
- During ageing, female mice fed a low protein diet during lactation have lower muscle weight.
- During ageing, female mice fed a low protein diet postnatally maintain their myofibre number.
- Male mice fed a low protein diet postnatally have lower body weight and muscle weight throughout their lifespan.
- Low protein diet affects myofibres size of *TA* muscle of male but not female mice at 3 months of age however this effect is lost during ageing.

## Introduction

Skeletal muscles develop in early embryological stages when the number of fibres is determined (Timson & Dudenhoeffer, 1990). Moreover, muscle mass and muscle fibre number depend on the proliferation and maturation of muscle precursor cells during embryological stages (Amthor et al., 1998). The late initiation of differentiation of precursor muscle cells leads to a bigger muscle mass at birth (Amthor et al., 1998). During the lifespan, skeletal muscles lose myofibres leading to a decrease in muscle mass that affects muscle function (Cruz-Jentoft et al., 2010). In humans, the loss of muscle fibres generally starts after the age of 30 and significantly decreases during ageing around the age of 70 leading to loss of muscle strength and frailty (Marzetti et al., 2009). The loss of muscle fibres during ageing is known as sarcopenia (Lexell, 1995; Petermann-Rocha et al., 2022; Snijders et al., 2009). The underlying mechanisms leading to muscle loss and sarcopenia have not been fully identified. However, studies suggest that environmental factors such as a sedentary lifestyle and a poor diet play a role in inducing muscle loss (Cruz-Jentoft et al., 2010). It has been shown that poor maternal diet during early life stages negatively affects the growth and has a long-term effect on offspring (Beermann, 1983; McCoski et al., 2021; Rehfeldt et al., 2011).

Proteins are the building units of skeletal muscles, therefore dietary proteins were always considered as an essential nutrient to maintain growth, and repair of muscle mass and function (Carbone & Pasiakos, 2019; Uzman, 2003). Previous studies have shown that protein restriction during lactation inhibited the development of mammary glands, ultimately causing a decrease in the quantity and quality of the breast milk produced (Cambraia et al., 1997; Moretto et al., 2011; Zheng et al., 2021). A protein restricted diet during lactation led to a 12% decrease in the protein content of the milk (Derrickson & Lowas, 2007). This inadequacy of milk supply leaves the new-borns malnourished, causing delays in their growth and a decrease in their overall body weight, which can have short-term and long-term impacts on offspring (Cambraia et al., 1997; Moretto et al., 2011; Zheng et al., 2021). This decrease in body weight and delayed growth of offspring was associated with several metabolic diseases such as type 2 diabetes and also with sarcopenia (Alexander et al., 2014; Beermann, 1983; da Silva Aragão et al., 2014; Loche & Ozanne, 2016; Ortega et al., 2013; Patel et al., 2012). Furthermore, studies have reported that short-term protein restriction during lactation (i.e. normalised protein intake post-weaning) in humans and rodents led to muscle wasting, and reduction in muscle force, muscle weight, and muscle fibre diameter (da Silva Aragão et al., 2014; Giakoumaki et al., 2022; Toscano et al., 2008; Wu, 2016). In addition, previous studies showed a correlation between induced slow growth rate of offspring caused by lactational low protein diet (LPD) and longevity (Chen et al., 2009; Heppolette et al., 2016; Lamming et al., 2015).

Lactational LPD can have a different effect on skeletal muscle of male and female offspring. Systems regulating energy metabolism differ between males and females leading to different responses to maternal diet. Other factors such as hormones, adipokines, placenta, and epigenetics also play a role in the sexual dimorphism in maternal diet studies. For example, oestrogen is a key hormone increasing the number of mitochondria while testosterone reduces it (Capllonch-Amer et al., 2014). The different hormonal effect explains the differences in the lifespan and response of males and females to maternal diet (Dearden et al., 2018).

The long-term effects of LPD on skeletal muscle during ageing have not been previously established. The aim of this study was therefore to determine the life-long effects of lactational low protein diet on muscle mass and function of male and female progenies. We hypothesised that a protein-deficient diet during lactation promotes muscle loss and would lead to an early onset of sarcopenia.

## Materials and Methods

### 1. Animals and experimental groups

The aim of this experiment was to study the effect of maternal LPD during lactation on the development and ageing of mice male and female progenies. C57BL/6 Thy1-YFP16 mice (The Jackson Laboratory; *Thy-1 YFP-16*, Stock# 003709) were housed in the Biomedical Service Unit (BSU) at the University of Liverpool and subjected to 12-hour light-12-hour dark cycle. All experiments described here were ethically approved by the University of Liverpool Animal Welfare and Ethical Review Body (AWERB) and performed in accordance with UK Home Office guidelines under the UK Animals (Scientific Procedures) Act 1986.

Female mice were fed either a normal (N) protein diet (20% Crude Protein W/W ISO’ (P), Code 829206, Special Diet Services, Essex, UK) or a low (L) protein diet (8% Crude Protein W/W ISO’ (P), Code 829202; Special Diet Services, Essex, UK) were mated with male mice fed a normal protein diet. The females were fed their respective diets from 2 weeks before mating, and throughout mating, birth and lactation. The normal and low protein diets are isocaloric.

New-born pups born from a mother fed NPD were cross-fostered and lactated by a dam fed either a NPD or a LPD leading to two different groups of mice: NN and NL. The number of suckling pups was kept similar for both NPD and LPD lactating mouse dams. After 21 days, weanling mice were fed either a NPD or a LPD throughout their lifespan leading to three different groups: NNN, NLN, and NLL. NNN group represents the control group, NLN mice were fed a LPD only during lactation and NLL mice were fed a LPD during their lifespan. Healthy mice were culled through a rising concentration of carbon dioxide (CO_2_) at 21 days (21d), 3 months (3M), 18 months (18M) and 24 months (24M) of age. Body weight was measured immediately after culling. NN and NNN groups served as controls for 21d and 3M/18M/24M, respectively (Fig. 3).

**Figure 1.**
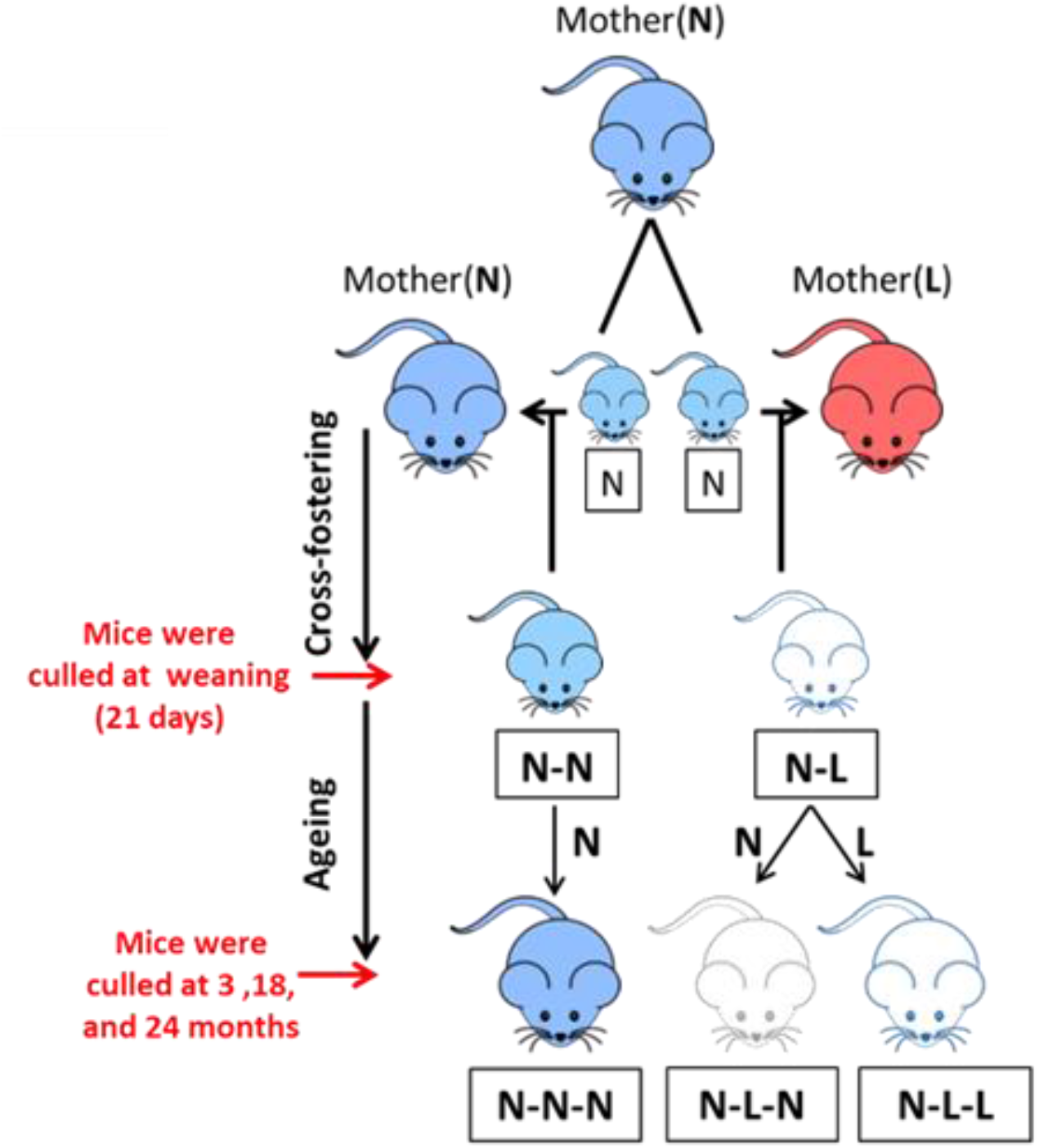
Experimental design (N) 20% diet (L) 8% diet. A) Group N: Control mice produced from a mouse dam fed a normal protein diet. Group N-N: Mice produced from a mouse dam maintained on a normal protein diet and fed postnatally by a mouse dam maintained on a normal protein diet. Culled and analysed at weaning (21 days). Group N-L: Mice produced from a mouse dam maintained on a normal protein diet but fed postnatally by a mouse dam maintained on a low protein diet. Culled and analysed at weaning (21 days). Group N-N-N: Control mice produced from a mouse dam fed on normal protein chow, fed postnatally by a mouse dam maintained on a normal protein diet until weaning and maintained on a normal protein diet. Culled and analysed at 3 months, 18 months and 24 months. Group N-L-N: Control mice produced from a mouse dam fed on normal protein diet, fed postnatally by a mouse dam maintained on a low protein diet until weaning and maintained on a normal protein diet. Culled and analysed at 3 months, 18 months and 24 months. Group N-L-L: Control mice produced from a mouse dam fed on normal protein diet, fed postnatally by a mouse dam maintained on a low protein diet until weaning and maintained on a low protein diet. Mice were culled and analysed at 21 days, 3 months, 18 months and 24 months.

### 2. Grip strength

To analyse the effect of maternal LPD on the physical performance of progenies during ageing, grips strength was measured. Grip strength of 18-month-old mice was assessed monthly using the four-limb hanging test. Mice were placed on a grid to acclimatise, and then the grid was inverted while the mice gripped using 4 limbs. The hanging time was recorded from the moment the grid was inverted until the mice lost their grip and fell onto a 5 to 7cm soft bedding. The mice were left to rest then the test was repeated (Carlson et al., 2011). The data was analysed as maximum hanging time and holding impulse (hanging time x body weight) taking into consideration the body weight of the mice.

### 3. Histological analysis

The purpose of this experiment was to study the histological effect of lactational LPD on the *Tibialis anterior* (TA) muscles of male and female progenies. *TA* were dissected, weighed, processed, and embedded in cryomatrix (Fisher Scientific, Loughborough, UK) then immersed in iso-pentane (Fisher Scientific, Loughborough, UK) cooled in liquid nitrogen (LN2) for 30 seconds and stored at -80°C.

Prior to sectioning samples were left in the cryostat (Leica 1900) that was set at -20°C for 30 min for acclimatisation. Sections of 10 μm thickness were collected on superfrost plus adhesion slides (Ct# J1800AMNZ, ThermoFisher Scientific, USA). The sections were stored at -20°C for staining.

Before staining, muscle sections were left at room temperature for 30 min to air-dry. The sections were washed with ice-cold phosphate buffered saline (PBS) for 2 min to wash away the cryomatrix, then fixed with ice-cold methanol (MetOH; Cat# 34860, Sigma Aldrich, Dorset, UK) for 5 min at room temperature. The fixed sections were washed thrice with PBS-Tween 20 (0.04% v/v; Sigma Aldrich, Dorset, UK) for 5 min each. Muscle sections were incubated in 1:1000 Wheat germ agglutinin (WGA– fluorescein 5 μg/mL; Cat# FL-1021-5, Vector Laboratories Ltd., Peterborough, UK) for 10 min followed by three washes of PBS-T for 5 min each. The sections were washed with H_2_O and left to dry for a few hours. Finally, the sections were mounted with hard-set DAPI (4′, 6-diamidino-2-phenylindole –hard set; Cat# H-1500, Vector Laboratories Ltd., Peterborough, UK). The slides were left overnight in the dark at 4°C. Before scanning, the slides were left at room temperature to dry, and then they were cleaned with 70% ethanol. The sections were scanned with a Z1 digital slide scanner (Carl Zeiss Microscopy, NY, USA). Images were acquired using 20x objective. Slides were kept in the dark at all times. Finally, images were analysed by using a fully automated system: MyoVision (University of Kentucky, USA) to measure the cross-sectional area (CSA) of the myofibres and count the number of fibres per section (Fig 2).

**Figure 2.**
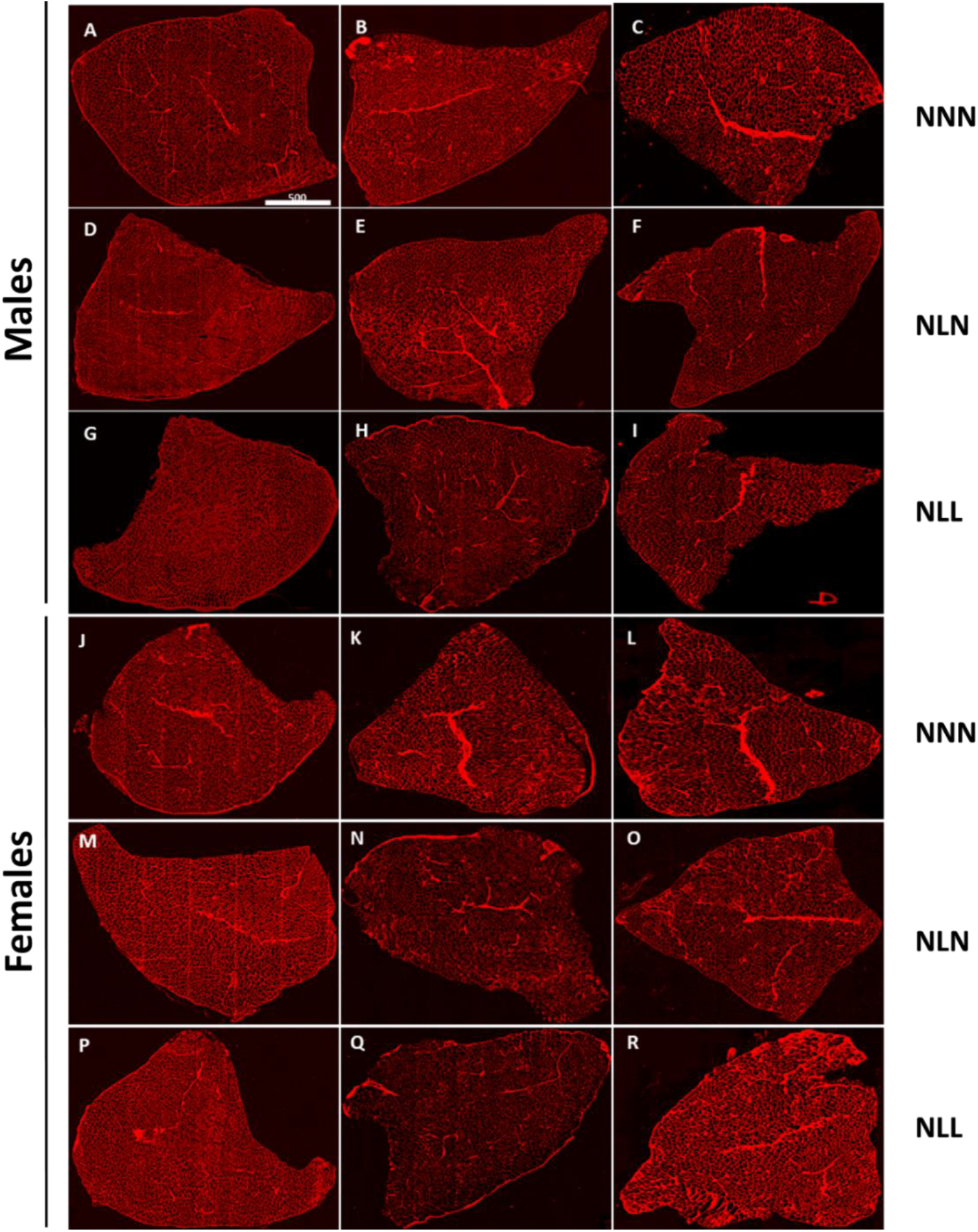
Representative cross-sections of TA muscle stained with WGA and imaged with Z1 digital slide scanner of male (A-I) and female (J-R). Cross section of TA of NNN male at 3 months (A), 18 months (B) and 24 months (C) of age. Cross-section of TA of NLN male at 3 months (D), 18 months (E) and 24 months (F) of age. Cross section of TA of NLL male at 3 months (G), 18 months (H) and 24 months (I) of age. Cross section of TA of NNN female at 3 months (J), 18 months (K) and 24 months (L) of age. Cross section of TA of NLN female at 3 months (M), 18 months (N) and 24 months (O) of age. Cross section of TA of NLL female at 3 months (P), 18 months (Q) and 24 months (R) of age. 20x magnification. Scale: 500 μm.

### 4. Statistical analysis

The data was analysed using GraphPad Prism 8 (version 8.02) and presented as mean ± standard error of the mean (S.E.M). Student-t test was used to compare NN and NL groups, and One-way ANOVA was used to compare NNN, NLN and NLL groups followed by Tukey’s post-hoc analysis comparing all groups. The data was considered statistically significant with a p-value less than 0.05.

## Results

### 1. Lifelong effects of lactational LPD on body weight and *TA* muscle weight of male and female mice

Lactational low protein diet significantly affected the body weight and muscle weight of male and female mice. Protein-deficient diet during lactation led to a decrease in the weight of both male and female mice at weaning age (NL-21 days) (Fig. 3 A and D). The male mice fed a LPD postweaning (NLL) maintained a significantly lower body weight at 3 months and 18 months of age compared to NNN and NLN mice. At 24 months, NLL male mice had a non-significant lower body weight because NNN and NLN mice significantly lost weight. The male mice fed a NPD post-weaning catch up with NNN mice at 3 months of age, but they show accelerated weight loss during ageing in comparison to NNN mice (Fig. 3A). NLN female mice also catch up with NNN female mice at 3 months of age. However, NLL female mice catch up by 18 months of age (Fig. 3D).

**Figure 3.**
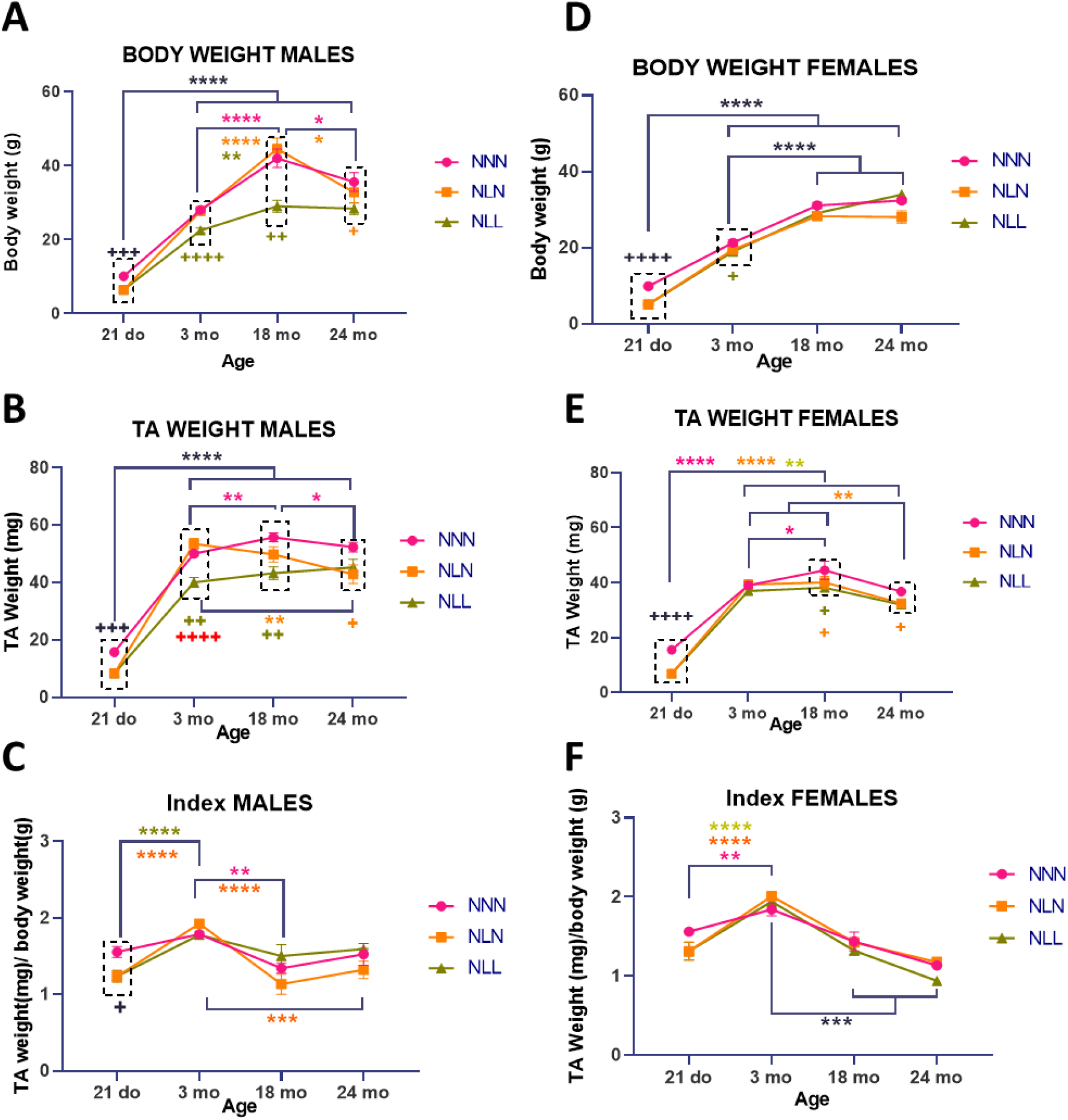
The effect of LPD during lactation on body and muscle weight, and index of male (A-C) and female (D-F) progenies. The effect of LPD during lactation on body weight of male mice throughout lifespan of NNN, NLN and NLL (A). The effect of LPD during lactation on TA weight of male mice throughout lifespan of NNN, NLN and NLL (B). The effect of LPD during lactation on index of male mice throughout lifespan of NNN, NLN and NLL (C). The effect of LPD during lactation on body weight of female mice throughout lifespan of NNN, NLN and NLL (D). The effect of LPD during lactation on TA weight of female mice throughout lifespan of NNN, NLN and NLL (E). The effect of LPD during lactation on index of female mice throughout lifespan of NNN, NLN and NLL (F). Data is presented as mean ± SEM, 21 days old (n=6-10), 3 months old (n=8-11), 18 months old (n=3-8), 24 months old (n=2-9), One-Way ANOVA followed by post-hoc Tukey’s test. * p≤0.05, ** p≤0.01, and **** p≤0.0001 representing NNN group, ** p≤0.01, and **** p≤0.0001 representing NLN group, ** p≤0.01, and **** p≤0.0001 representing NLL group, ^+^ p≤0.05, ^+++^ p≤0.001, ^++++^ p≤0.0001 to compare NN and NL at 21 days of age, ^+^ p≤0.05, ^++^ p≤0.01, ^++++^ p≤0.0001 to compare NNN and NLL at 3, 18 and 24 months of age, ^+^ p≤0.05, ^++^ p≤0.01, to compare NNN and NLN at 3, 18 and 24 months of age.

*TA* weight in male mice showed a similar pattern as the body weight (Fig. 3 B). NL male mice had a lower *TA* weight at 21 days of age, when fed a LPD postweaning they kept their low *TA* weight. At 24 months, the weight difference between NLL and NNN was not significant because *TA* weight of NNN male mice significantly decreased. *TA* weight of NLN male mice caught up with NNN mice at 3 months of age, however they showed an accelerated loss of muscle weight during ageing. Suggesting it is better to stay on a LPD lifelong rather than a short period of time. NL male mice had a lower ratio of muscle weight to body weight (index) at 21 days of age (Fig. 3C). However, this difference was no longer present at 3 months indicating that the loss of muscle weight was in proportion to body weight. During ageing, index of all male mice decreased. Although not significant, NLN male mice had lower index than NNN and NLL mice potentially indicating a higher loss of muscle mass.

The *TA* weight of female mice showed a different pattern to the males. At 21 days of age, the *TA* weight was lower in the NL group (Fig. 3 E), however both NLN and NLL mice caught up with NNN mice at 3 months of age then showed an accelerated muscle weight loss during ageing (Fig. 3 E). This data suggests that short-term and long-term LPD negatively affect muscle weight of female mice during ageing. Moreover, NL female mice have lower muscle to body weight index at 21 days, however similar to males, both groups show similar rate of muscle wasting in adulthood when adjusted for body weight (Fig. 3F).

### 2. The lifelong effect of LPD during lactation on myofibres of *TA* muscle in male and female mice

Histological analysis of *TA* muscle showed that the number of myofibres in male offspring was not different among the groups at 3, 18 or 24 months of age (Fig. 4 A). NNN and NLN male mice lost fibres during ageing specifically at 18 months of age, unlike NLL mice that had a smaller fibre number at 3 months and showed a faster decline rate (Fig. 4 A). The CSA of the myofibres was significantly lower in NLL male mice in comparison to NLN and NNN mice at 3 months of age (Fig. 4 B). During ageing, the CSA was not different among the groups which can be due to the increase in the CSA of myofibres of NLL mice at 18 months of age. NNN and NLN were similar in terms of myofibre number and CSA. Even more the distributions of myofibres of NNN and NLN were very similar, while NLL had a curve skewed to the left indicating that these mice had more fibres of smaller range at 3 months of age. During ageing, the distribution of myofibres of NLL was similar to NNN and NLN (Fig. 4 E, F and G) which correlates with the increase in the CSA of NLL myofibres at 18 months of age (Fig. 4 B).

**Figure 4.**
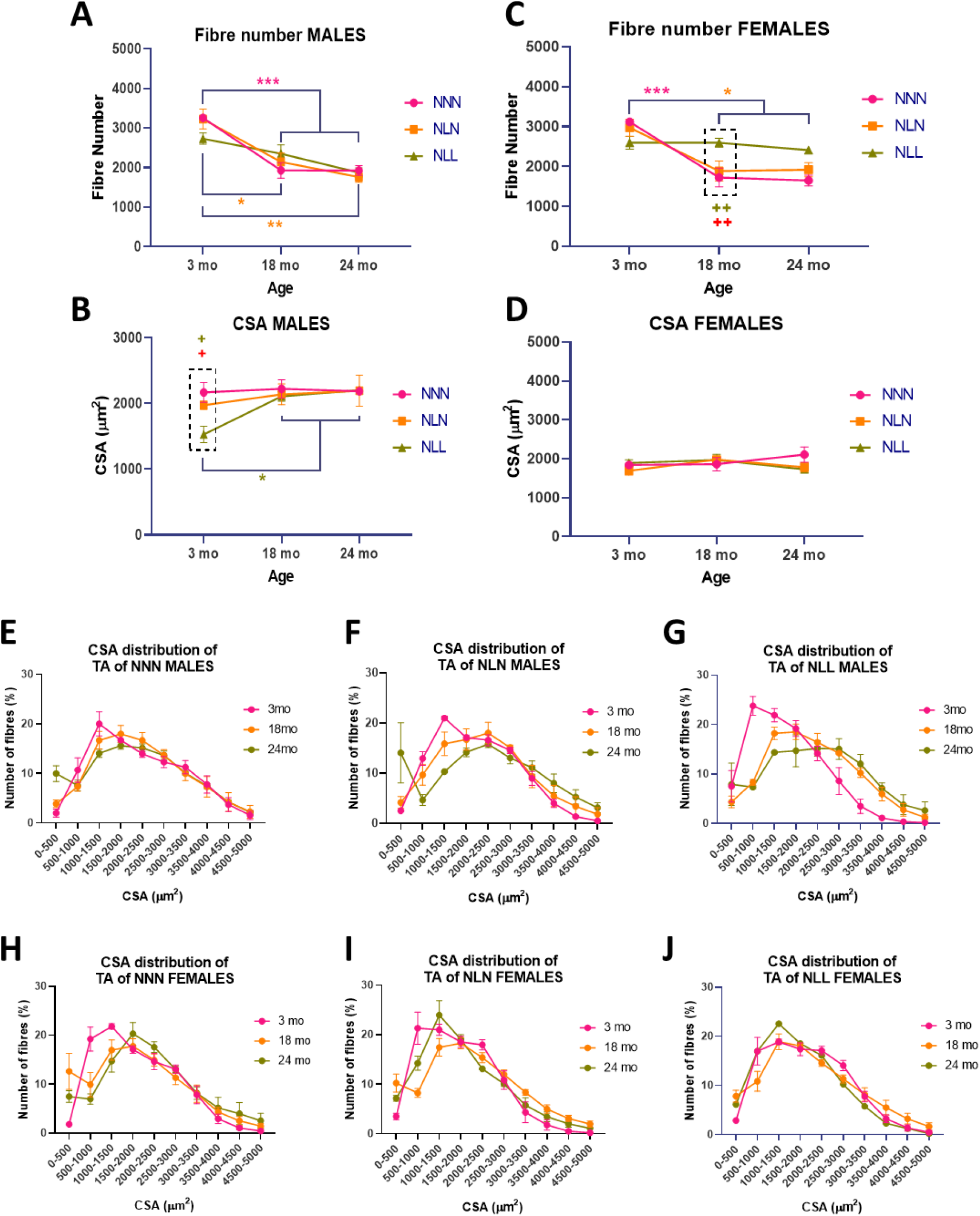
The effect of LPD during lactation on myofibres of TA muscle of male (A-B, E-F) and female (C-D, H-J) progenies. The effect of LPD during lactation on total fibre number of TA muscle of males (A) and females (C) throughout lifespan of NNN, NLN and NLL. The effect of LPD during lactation on CSA of myofibres of TA of males (B) and females (D) of NNN, NLN and NLL. The distribution of myofibres of NNN, NLN and NLL of males (E-G) and females (H-J). Data is presented as mean ± SEM, n=1-5. One-Way ANOVA followed by post-hoc Tukey’s test (A-D). Chi-square test and Two-Way ANOVA followed by post-hoc Tukey’s test (E-J). *** p≤0.001 representing NNN group, * p≤0.05, and ** p≤0.01representing NLN group, * p≤0.05 representing NLL group, ^+^ p≤0.05, and ^++^ p≤0.01 to compare NNN and NLL at 3 and 18 months of age, ^+^ p≤0.05, and ^++^ p≤0.01, to compare NLL and NLN at 3 and 18 months of age.

Similar to males, fibre number, CSA and fibre distribution in NLN females were similar to NNN (Fig. 4 C, D, and H-J). However, NLL female mice had a lower number of fibres at 3 months of age comparing to NNN and NLN and the number was preserved during ageing (Fig. 4 C). At 18 months of age, the number of myofibres of NNN and NLN mice decreased and had lower number than NLL female mice. LPD did not have a significant effect on the CSA of the myofibres or the distribution of the myofibres in females (Fig. 4 D and H-J). However, even though not significant both NLN and NLL had a lower CSA than NNN at 24 months of age. Only one NLL mouse survived until 24 months of age (Fig. 6 B) therefore no statistical analysis was possible.

### 3. The effect of LPD during lactation on grip strength of ageing male and female mice

Lactational LPD negatively affected the grip strength of male offspring. The maximum hanging time measured in this study was fluctuating throughout the measurements which can be due to the weight or behavioural factors while conducting the experiment. The hanging time of different groups in males and females was not significantly different between groups (Fig. 5 A-B and D-E). Holding impulse was calculated taking into consideration the body weight of the mice. Holding impulse of NLN male mice at 24 months normalised to 18 months was significantly lower than NNN mice (Fig. 5 C). Holding impulse of NLL mice was also lower than NNN mice; however, there was only one mouse in the group therefore statistical analysis was not possible. This suggests a higher decline rate in grip strength in NLN and NLL mice compared to NNN mice. The decline in NLN male mice might be due to their body weight as the change was only seen in holding impulse and not hanging time. Holding impulse of female mice was not different among the groups suggesting that body weight did not affect the grip strength of female mice.

**Figure 5.**
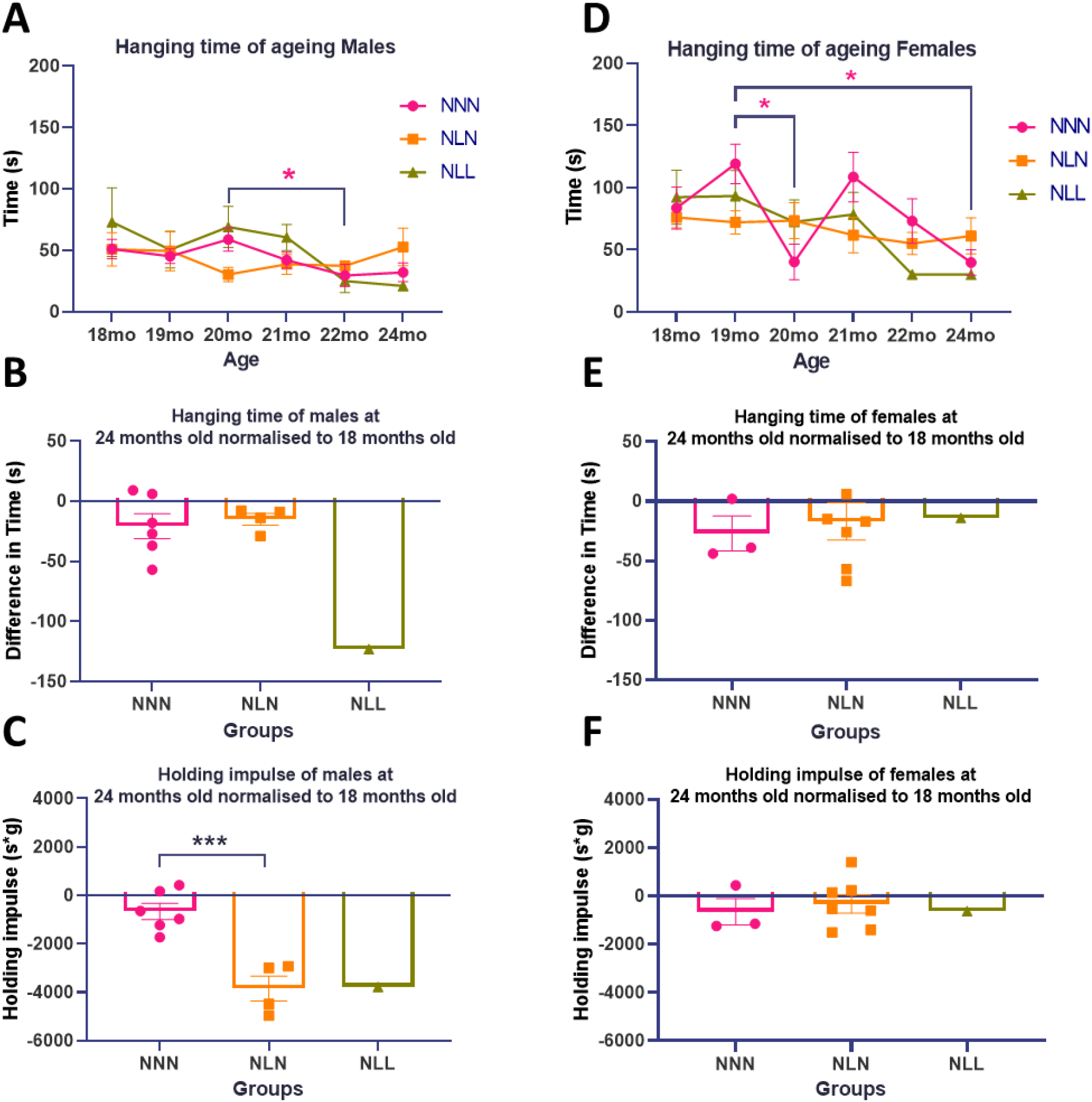
Grip strength of ageing male (A-C) and female (D-F) mice. Hanging time was measured as maximum hanging time and holding impulse (hanging time x body weight). Hanging time of ageing male (A) and female mice (D) (> 18 months of age) of NNN, NLN, and NLL groups measured monthly until 24 months of age. Hanging time of 24 months old male mice normalised to 18 months (B) and female mice (E). Holding impulse of 24 months old male mice normalised to 18 months (C) and female mice (F). Data is presented as mean ± SEM, NNN (n=5-10), NLN (n=3-6), NLL (n=1-4) at 18 months of age. Two-Way ANOVA followed by post-hoc Tukey’s test (A and D). * p≤0.05 in NNN (24mo data was excluded from the analysis because there was only one mouse in NLL group). Unpaired t-test to compare NNN and NLN while NLL was excluded from the analysis because there was only one sample (B-C and E-F-). *** p≤0.001.

### 4. LPD effect of lifespan

Postnatal LPD significantly decreased the lifespan of female mice (Fig. 6 B). Protein restriction during lactation followed by a normal diet after weaning (NLN) did not have any effect on the survival rate of either male or female (Fig. 6 A and B) offspring. The sample size of NLN mice at 18 months was 6 for males and 10 for females and only 2 death events happened until 24 months of age for males and females. However, postweaning LPD led to a non-significant decrease in the lifespan of male where only 2 death events occurred reducing the number of mice from 4 at 18 months to 2 at 24 months of age (Fig. 6 A). However, there was a significant decrease in survival rate of female progenies as there were 6 deaths events throughout the 6 months reducing the number of female mice from 7 at 18 months to 1 at 24 months of age (Fig. 6 B).

**Figure 6.**
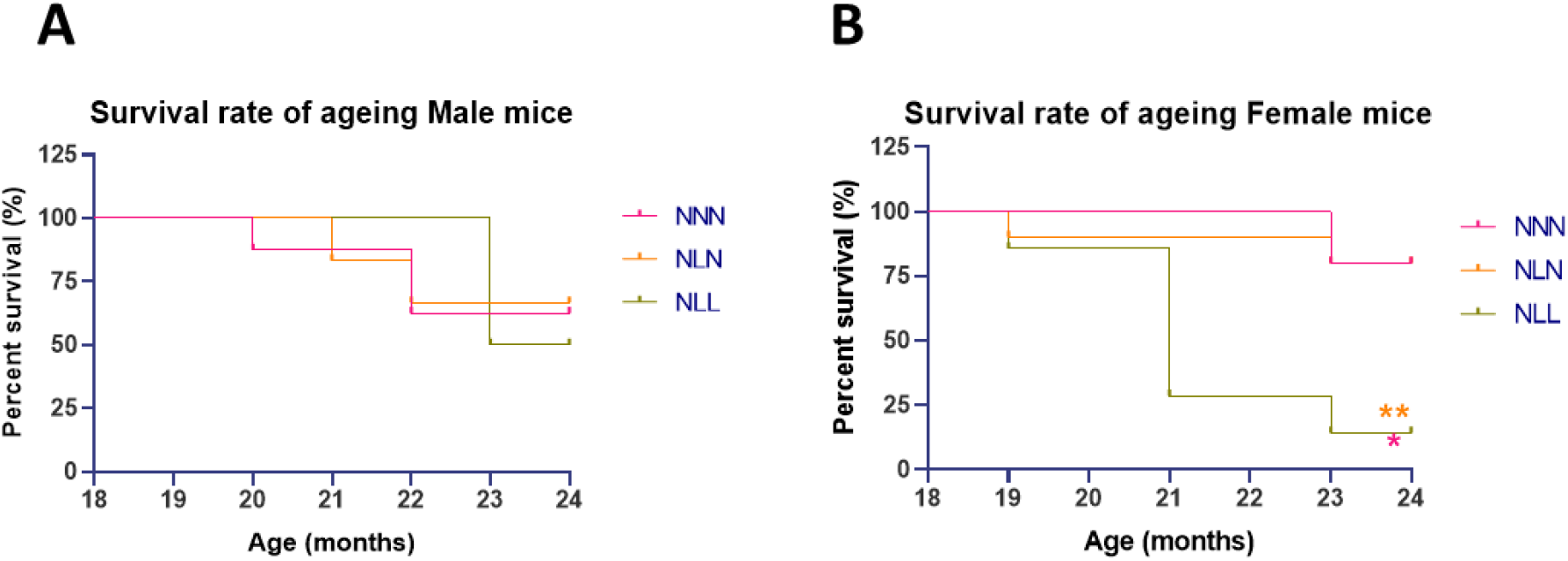
The effect of LPD during lactation on the survival rate of male (A) and female (B) progenies. NNN (n=8), NLN (n=6), NLL (n=4) at 18 months of age for males and NNN (n=5), NLN (n=10), NLL (n=**7**) at 18 months of age for females. Log-rank test, three groups analysis. * p≤0.05 for NLL compared to NNN, and ** p≤0.01 for NLL compared to NLN.

## Discussion

The aim of this investigation was to examine the impact of a maternal low protein diet during lactation on the development and ageing processes of skeletal muscle in male and female offspring. Various parameters including body weight, muscle weight, grip strength, and survival rate were evaluated at different life stages: weaning (21 days), early adulthood (3 months), early ageing (18 months), and late ageing (24 months). Furthermore, histological analyses of the *TA* muscle in the progenies were conducted. Additionally, we examined whether this dietary intervention, followed by either a sustained LPD or a transition to NPD, contributes to the early onset of sarcopenia.

LPD during lactation did not increase the lifespan of male and female progenies. In this study, the survival rate of aged female NLL mice was significantly lower compared to NLN and NNN mice. Although the survival rate of male NLL mice decreased by 50%, this decrease was not statistically significant when compared to NNN and NLN mice. On the other hand, the survival rate of NLN male and female mice was not affected by the diet and remained similar to that of NNN mice. Previous studies have suggested that maternal LPD and slow growth rate may extend the lifespan of rodent progenies (Chen et al., 2009; Guzmán et al., 2006; Heppolette et al., 2016; Ozanne & Hales, 2004). Both Heppolette et al. (2016) and Ozanne & Hales (2004) used the same diet as the one used in this study and focused on the effect of lactational low protein diet on male offspring. In all these studies the offspring remained smaller post-weaning in contrast to the current study where recuperation of weight was observed. Ozanne & Hales (2004) focused on the effect of slow growth rate postnatally due to LPD during lactation corrected post-weaning and found that it increases lifespan of male pups. Heppolette et al. (2016) correlated this increase in lifespan with delayed ageing of adaptive immune system. Guzmán et al., (2006) studied the effect of maternal low protein diet (10 %) on female progenies in rats and found that survival rate of NLN mice was not significantly lower than NNN mice even though they had only one mouse at 22 months of age. None of these studies had an NLL group in their studies. However, Carey et al., (2022), negatively correlated protein intake with the lifespan of male and female *Drosophila melanogaster* which contradicts with this study.

The impact of maternal LPD during lactation on body weight, muscle weight, grip strength, and survival rate, differed between male and female offspring. In this study, 21-day-old male and female mice that were fed a LPD during lactation (NL) exhibited a lower body weight compared to control mice (NN). This finding aligns with previous studies that have demonstrated a negative correlation between LPD during lactation and the body weight of offspring (Cambraia et al., 1997; Chen et al., 2009; Giakoumaki et al., 2022; Moretto et al., 2011). Interestingly, the body weight of male NL mice that were subsequently fed a NPD post-weaning (NLN) caught up with the body weight of NNN mice at 3 months of age and exhibited a growth pattern similar to the control group throughout the ageing process. Conversely, NLL male mice consistently maintained a lower body weight throughout their lifespan compared to both NNN and NLN mice. In the case of female mice, post-weaning LPD resulted in a different pattern compared to males. NLN and NLL female mice caught up with NNN mice in terms of body weight at 3 months and 18 months of age, respectively. This finding contrasts with the results of Heppolette et al., (2016), which showed that NLN male mice consistently maintained a lower body weight than NNN mice throughout their lifespan and did not show any data about females. Various studies have consistently demonstrated the negative effects of nutritional restriction on body weight and growth rate (Beermann, 1983; Victora et al., 2008). Catch-up growth following in utero growth restriction as a result of improvement of nutritional deficits during early life stages has been reported (Chen et al., 2009; Ozanne & Hales, 2004). LPD during adulthood has also been linked to a decrease in body weight due to metabolic alterations (Pezeshki et al., 2016), which provides an explanation for the lower body weight observed in NLL mice.

Postnatal LPD affects *TA* muscle of female and male progenies differently. In female offspring, those fed a LPD during lactation had lower *TA* muscle weight compared to control mice at 21 days of age. However, by 3 months of age, their *TA* weight caught-up to the control group regardless of the post-weaning diet. At 18 months of age, there was a decrease in *TA* weight, accompanied by a significant loss of muscle fibres in one group (NLN), resembling the pattern observed in control mice. The other group (NLL) did not experience fibre loss but showed a potential higher number of smaller range fibres. Additional factors such as fat infiltration, connective tissue changes, water retention, or fibre remodelling may contribute to the observed *TA* weight changes in NLL mice (Cho & Suh, 2016). In male offspring, LPD led to lower *TA* muscle weight in NL mice at 21 days of age, and although NLN mice caught up to the control group by 3 months of age, a significant decrease was observed at 24 months of age. NLL mice consistently had lower *TA* weight throughout their lifespan. Histological analysis indicated that LPD did not significantly affect the number of myofibres but had a negative impact on myofibre CSA in NLL mice at 3 months of age. Interestingly, the myofibre CSA increased during the ageing process, potentially due to the loss of smaller range fibres. The exact mechanism behind the *TA* weight loss in NLL male mice requires further investigation, potentially related to fat infiltration or connective tissues. Furthermore, it is worth to mention that the analysis at 24 months of age for males and females essentially is comparing the fittest progenies that survived up to that age.

LPD is known to inhibit protein synthesis and can lead to protein catabolism in skeletal muscle when associated with high carbohydrate diet (McGlory et al., 2019; Toscano et al., 2008). In humans, LPD was positively associated with muscle mass loss (Huang et al., 2016; Huh & Son, 2022). Previous studies showed lactational LPD in male progenies led to a decrease in skeletal muscle weight at 21 days of age and 3 months of age for NLN mice in comparison to control (Chen et al., 2009; Giakoumaki et al., 2022). In rats, LPD affected the triacylglycerol and glycogen levels in skeletal muscle but had no effect on skeletal muscle mass and fibres (Moretto et al., 2011). These studies showed different effects of LPD on skeletal muscle, but none of them studied the effect of lactational LPD during ageing nor the postnatal LPD. During ageing, skeletal muscle loses its mass (Deschenes, 2004) which is compatible with NNN and NLN mice in this study but not with NLL. This suggests that protein restriction during lactation normalised during adult life does not affect muscle mass. The *TA* weight of NLL mice both male and female were maintained during ageing, the mechanism behind this stability in *TA* weight needs to be further investigated. The long-term protein restriction in both male and female progenies prevented the muscle mass loss associated with ageing. Skeletal muscles are mainly made of myofibres that can affect muscle mass. However, skeletal muscle also contains fat and connective tissue that can also affect its mass. LPD was associated with protein degradation and loss of myofibres due to metabolic changes caused by protein deficiency during adulthood (Pezeshki et al., 2016). Surprisingly, in this study the number of myofibres of *TA* muscle of mice fed a LPD postnatally (NLL) were maintained during ageing. LPD was also associated with a switch of type II to type I myofibres which are smaller than type II (Toscano et al., 2008). This switch in myofibre types is also associated with ageing (Nakazato & Song, 2008; Schiaffino & Reggiani, 2011; Toscano et al., 2008). Unfortunately, in this study the fibre type of the *TA* muscle was not assessed however the data showed that LPD affects CSA in NLL male mice at 3 months which can be associated with myofibre remodelling. In females, even though not significant, NLL mice had a lower CSA however only one mouse survived until 24 months of age. Male and female mice are known to have different skeletal muscle composition, contractility, and metabolism (Haizlip et al., 2015). Sex hormones also affect skeletal muscle physiology. For example, oestrogen which is a female hormone is associated with skeletal muscle maintenance and high muscle regeneration while testosterone a male hormone is associated with muscle growth (Maher et al., 2010). Zambrano et al., (2005), showed that NLN rats had an increased testosterone level at early stages of life, but after 6 months testosterone level decreased which might explain the decrease in *TA* muscle weight in NLN during ageing and the catch-up growth of *TA* muscle at 3 months of age. Maternal LPD during lactation was associated with a decrease in oestrogen level which delays the development of reproductive organs and induce early ageing of reproductive system (da Silva Faria et al., 2010; Guzmán et al., 2006). The low oestrogen level which was associated with lactational LPD was also associated with muscle loss especially during menopause in females which contradicts with the data in this study. Further investigation is needed to understand the underlying mechanism of muscle loss and maintenance in male and female mice.

Lactational LPD led to a decrease in the holding impulse of old male mice but did not have an effect on the grip strength of female mice. The decline in muscle strength along with muscle mass is characteristic of sarcopenia (Metter et al., 1997). The decrease in muscle activity was associated with changes in muscle physiology. For example, myofibre size and myofibre type affect physical activity and muscle strength. Protein restriction was also associated with a decrease in physical performance (Beermann, 1983; da Silva Aragão et al., 2014; Granic et al., 2018; Toscano et al., 2008).

## Conclusion

In conclusion, the findings of this study highlight the impact of a lactational LPD on the development and ageing of male and female offspring. Firstly, it was observed that mice fed a LPD during lactation had a lower body weight compared to control mice. However, NLN male mice eventually caught up with their counterparts fed a normal protein diet, indicating a compensatory growth response. Unlike males, female NLN and NLL mice eventually caught-up with NNN mice. Furthermore, the effects of the LPD on the *TA* muscle differed between male and female mice. In male mice, the LPD primarily affected the size of myofibres of the *TA* muscle. On the other hand, in female mice, the LPD predominantly influenced the number of myofibres in the *TA* muscle. Notably, a significant finding was the impact of postnatal LPD on the survival rate of female progenies. These results shed light on the importance of proper nutrition during lactation and its potential implications on the ageing of offspring. Additional research is needed to understand the underlying mechanisms involved in the loss or preservation of muscle mass during ageing, as well as the distinct mechanisms operating in male and female offspring.

## Ethics approval

All experimental protocols were performed according to the UK Animals (Scientific Procedures) Act 1986 regulations and obtained ethical approval from the University of Liverpool Animal Welfare Ethical Review Board (AWERB). All animal procedures were performed under a valid Home Office Project Licence.

## Availability of data and materials

Not applicable

## Informed consent statement

Not applicable.

## Author contributions

I.K., A.V. and K.G.W. designed the experiments with input from S.E.O.; M.A. and I.K performed the grip strength, dissection and tissue processing. M.A performed the staining, analysed the data and drafted the manuscript. A.E.A., E.U., and I.M.K., provided help with the analyses and interpretation of the data. All authors have read and agreed to the published version of the manuscript.

## Acknowledgements

We thank the Biomedical Services Unit at the University of Liverpool. The authors also thank the Rank Prize for their financial support.

